# Transverse sinus injections: A novel method for whole-brain vector-driven gene delivery

**DOI:** 10.1101/579730

**Authors:** Ali S. Hamodi, Aude Martinez Sabino, N. Dalton Fitzgerald, Michael C. Crair

## Abstract

A major challenge in neuroscience is convenient whole-brain delivery of transgenes, especially early in postnatal development. Recent advances demonstrate whole-brain gene delivery by retro-orbital injection of virus, but slow and sparse expression, and the large injection volumes required, precludes early developmental studies. We developed a novel method for simple, fast and efficient gene delivery across the central nervous system in neonates as early as P4 and persisting into adulthood. The method employs transverse sinus injections of 2-4μL of AAV9 at P0. Here, we describe how to use this method to label and/or genetically manipulate cells in the neonatal rat and mouse brain. This protocol is fast, easy, can be readily adopted by any laboratory, and utilizes the widely available AAV9 capsid. The procedure outlined here is adaptable for diverse experimental applications ranging from biochemistry, anatomical and functional mapping, to gene expression, silencing, and editing.

## Introduction

Recombinant adeno-associated viruses (AAVs) are commonly used vectors for *in vivo* gene delivery [1], and recently published work demonstrates whole-brain gene delivery by retro-orbital injection of AAV9 and other engineered AAV variants [2]. However, the onset of robust systemic expression of genes through AAVs traditionally occurs several weeks after the time of injection [2]. This presents a particular challenge for experiments requiring gene expression early in postnatal development. In addition, current gene-delivery methods, such as retro-orbital injections, are difficult and particularly disruptive in early postnatal mice. Other alternatives, such as intravenous administration of virus through the tail or temporal vein, require high injection volumes (100μL) that are expensive and disruptive for P0-P1 pups, and still produce inefficient transduction (15-18%) of target cells [1]. Other alternatives, such as targeted gene expression through the creation of transgenic mice, are also limited by the complexity of interbreeding, which often necessitates the crossing of two or more strains to drive conditional expression in the desired cell types [3,4].

To remedy this, we developed an easy and efficient method of transgene delivery through the transverse sinus in neonatal mice, which we refer to as ‘n-SIM’. The method targets the transverse sinus, which is easily accessible along the posterior edge of the forebrain and convenient for virus injection in neonatal mice when the skin and skull remain quite thin. AAV9 is very well suited for this approach given the proclivity of AAV9 to cross the blood-brain barrier (BBB), especially before it is fully developed in neonates [1]. The position and proximity of the transverse sinus to the brain also makes it particularly well suited for successful gene delivery to the brain. Indeed, the injection of as little as 2-4μL of AAV9 (1×10^13^ vg/mL) into the transverse sinuses of neonatal mice results in robust and widespread gene delivery to the brain. With virus injection at P0, we observe dense labeling in cortex, thalamus, midbrain, and hippocampus as early as P4, which persists into adulthood. Sinus injections with AAV9 were successfully tested in both mice and rats but are likely suitable for any mammalian species.

This method enables the targeting of distinct cell populations at early stages of development and enables the delivery of multiple viral constructs at the same time across the whole brain. This will dramatically accelerate the application of novel molecular technologies without the need to generate costly transgenic strains or complicated crosses. Sinus injections also circumvent the caveats of direct injections into the brain parenchyma, which can cause tissue damage and variable gene expression. This is especially important for neurodevelopmental studies, but the approach is applicable to older animals as well, as the expression persists in adults. Our method provides an easy and fast way (10 min per pup) to express a wide variety of transgenes across the extent of the brain using the easily accessible AAV9 capsid. Finally, sinus injected animals show no deleterious health effects, either from the injections or as a result of high expression levels typically associated with early embryonic expression in transgenic mice [4]. We describe below in detail how our method is performed and demonstrate the ability to carry out experiments in mice and rats to answer fundamental questions in neuroscience that were previously impractical or impossible.

## Methods

All experimental procedures are in accordance with National Institutes of Health guidelines and approved by Yale Institutional Animal Care and Use Committees. Animals are treated in compliance with the U.S. Department of Health and Human Services and Yale University School of Medicine. To validate our method and test its applicability, we perform two-photon and widefield calcium imaging *in vivo* experiments on mice we’ve injected using our method, immunohistochemistry, cross-method validation experiments, and control experiments.

### Mice

To label the different interneuron subtypes, we used Nkx2.1-cre, SOM-cre, VIP-cre, LSL-tdTomato (The Jackson Laboratory (JAX) strains 008661, 018973, 010908, 007909, ME, USA) mice. All mice were housed on a 12-hour light/dark cycle with food and water available *ad libitum*. For histological validation of sinus injections, we used C57BL6/J mice (JAX 000664). For rat experiments, we used Long-Evans rats (strain code: 006, Charles River, MA, USA).

### Neonatal transverse sinus injection method (n-SIM)

P0-P1 pups were removed from the cages and placed on a warm pad. Each pup rested on ice for anesthetization for 2-3 minutes before being transferred to a cold metal plate where we performed the injection. A light microscope was used to visualize the transverse sinuses (located on the dorsal surface of the mouse head, Supplementary figure 1A,B). Next, sterilized fine scissors (Fine Science Tools, CA, USA) were used to make two small cuts (~2mm) in the skin above each transverse sinus (Supplementary figure 1C).

To inject the virus, we used capillary glass tubes (3.5” #3-000-203-G/X, Drummond Scientific Co, PA, USA), pulled using a P-97 pipette puller (Sutter Instruments, CA, USA) to produce fine tips with high resistance. The sharp pipettes were filled with mineral oil (M3516, Sigma-Aldrich, NY, USA) then attached to a Nanoject III (Drummond Scientific Co). Most of the mineral oil was ejected using the Nanoject. Next, the vector solution was drawn into the pipette. For accurate movement of the Nanoject-attached-pipette, we used MP-285 micromanipulator (Sutter Instruments).

The pipette was gently lowered through the skull and into the sinus until the tip of the pipette was observed (using the light microscope) to break through the sinus vessel wall. The pipette tip was retracted until it was 300-400μm below the surface of the skull, such that the tip resided within the lumen of the sinus. With no delay, 2 or 4μL of virus was injected at a rate of 20nL per second. Following a 5 second delay, the pipette was retracted, and the same loading and injection procedure was repeated targeting the opposite hemisphere. Virus injections per mouse pup were in a total volume of 4μL of AAV9-syn-GCaMP6s (Addgene, MA, USA) with a titer of 1Χ10^13^ vg/mL. For RCaMP experiments, we co-injected a total volume of 4μL of 1Χ10^13^ vg/mL of AAV9-CAG-flex-GCaMP6s (Addgene) and AAV9-syn-jRCaMP1b (Addgene) (1:1 ratio) at P1. A successful injection was verified by visualizing viral solution flow in the blood stream and blanching of the sinus (Supplementary figure 1C’,C”). After the injection, the skin was folded back, and a small amount of VetBond glue was applied to the cut. The pup was then returned to the warm pad. After the whole litter was injected, the pups were returned to their home cage and gently rubbed with bedding to prevent rejection by the mother.

### Perfusion

After allowing time for expression (4 days minimum), neonatal mice were anesthetized with Ketamine/Xylazine/Acerpromazine mix (37.5 mg/mL, 1.9 mg/mL, and 0.37 mg/mL, respectively). The anesthetic dose for this combination cocktail was 2.0-3.0 mL/kg, and it was administered via intraperitoneal (IP) injection. Next, mice were transcardially perfused with phosphate buffer saline (PBS) at room temperature and then with freshly prepared, ice-cold 4% paraformaldehyde (PFA) solution at 5, 9, 13, or 20 days post-injection. Brains were then extracted, and immersion fixed in 4% PFA overnight, then rinsed with PBS.

### Comparison of AAV9 n-SIM to other methods

To compare the efficacy of AAV9 n-SIM at P1 relative to other AAV serotypes, we injected the same volume (4μL) of either AAV1-syn-GCaMP6s (titer: 1Χ10^13^ vg/mL, Addgene), AAV5-syn-GCaMP6s (titer: 1Χ10^13^ vg/mL, Addgene), or AAV.PHPeB-syn-GCaMP6s (titer: 1Χ10^13^ vg/mL, Gradinaru Lab). All injection conditions were the same as described above. To compare n-SIM to temporal vein injections, the former was performed using the same anesthetization procedure described above. Next, we made a small cut to expose the temporal vein, and we used a Nanjoject to inject the virus into the vein. After injecting the temporal veins on each side, the cuts were sealed with Vetbond (Vetbond, 3M, MN, USA) and the pups were returned to their home cage.

### Histological processing, immunohistochemistry and imaging

To validate and quantify the levels of expression after n-SIM, neonatal brains were sectioned into 150μm coronal or sagittal slices using a Leica VT1000S vibratome (Leica, IL, USA). Slices were transferred into 0.04% Triton solution, then blocked overnight with 10% goat serum at 4°C. After blocking, primary antibodies were diluted in the blocking solution (1:500), and slices were incubated in the primary antibody solution for 5 days at 4°C. The primary antibodies used were rabbit anti-GFP conjugated to AF488 (Millipore), and mouse anti-NeuN (Millipore). Next, slices were washed with PBS (3Χ15 min), and incubated in secondary antibody diluted in blocking solution (1:500) overnight at 4°C. The secondary antibody used was goat anti-mouse AF555 (Millipore). Next, slices were washed with PBS (3Χ15 min), incubated in anti-DAPI antibody diluted in PBS (1:1000) for 15 min, washed with PBS (3Χ15 min), and mounted on glass slides using Flouromount G (Vectashield, Vector Laboratory, CA, USA) then coverslipped. For display purposes and assessment of overall brightness, slice images were captured using Zeiss Apotome microscope (Zeiss, Oberkochen, Germany) using exposure time and contrast settings described in the figure legends. For quantification, slices were imaged using Zeiss laser scanning confocal microscope (LSM 800) equipped with EC Plan-Neofluar 10Χ/0.3 (Working Distance=5.2mm, Zeiss) and Plan-Apochromat 20Χ/0.8 (Working Distance= 0.55mm, Zeiss) objectives to determine co-localization between GCaMP6s+ and NeuN+ signals. Signal quantification was done using ImageJ software. Briefly, images were binarized, regions of interest (ROIs), were selected, and the percent overlap between GCaMP6s+ and NeuN+ cells was manually counted twice by multiple blinded observers to minimize bias.

### *In vivo* imaging

#### Surgical preparation

To prepare the animals for functional imaging, mice and rats were anesthetized using 1-2% isoflurane and maintained at 37°C using a water heating pad for the duration of the surgery. The scalp was cleaned with povidone-iodine solution, then topical lidocaine applied and Maloxicam (0.3 mg/kg) administered IP. The skin and fascia layers above the skull were removed to expose the entire dorsal surface of the skull from the posterior edge of the nasal bone to the middle of the interparietal bone, and laterally between the temporal muscles. The skull was thoroughly cleaned with saline, and the edges of the skin incision secured to the skull using Vetbond glue.

To head-fix the animal, we used a custom headpost which consists of two screws (0/80 × 3/16 MS24693-C420 Phillips Flat Head 100° 18/8 Stainless Steel Machine Screws, Mutual Screw & Supply, NJ, USA) placed upside down on the skull (i.e. base first) and secured onto the interparietal and nasal bone with Vetbond, and transparent dental cement (Metabond, Parkell, Inc., NY, USA). To reduce motion and exposure of bone to air, and a thin layer of dental cement was applied to all exposed skull. Once the dental cement was dried, it was covered with a thin layer of cyanoacrylate (Maxi-Cure, Bob Smith Industries, CA, USA) to provide a smooth surface for imaging. The head-screws were threaded through the holes of a custom-made metal bar and secured with metal nuts. For two-photon imaging, we performed a 3mm diameter cranial window over visual cortex of the right hemisphere using a dental drill (Ram Products, Inc., NJ, USA), and the cranial window was covered with a double coverslip (Small round cover glass, #1 thickness, 3mm + Small round cover glass, #1 thickness, 5mm, Warner Instruments LLC, CT, USA) and sealed using Maxi-Cure. All mice were allowed to recover from head-post surgery for a minimum of three hours before imaging.

#### Widefield and two-photon calcium imaging

To validate functional expression of GCaMP across the cortical mantle, widefield calcium imaging was performed using a Zeiss Axiozoom V.16 with PlanNeoFluar Z 1Χ, 0.25 NA objective. Epifluorescence excitation was performed using a LED source (X-cite TURBO XLED, MA, USA) with 450-495nm illumination for GCaMP6s and 540-600nm for jRCaMP1b through a Fitc/Tritc filter cube (Chroma, 59022, VT, USA). Epifluorescence emissions were filtered with Semrock FF01-425/45-25, Semrock FF01-624/40-25, and Semrock FF01-593/LP-25 (Semrock Inc., NY, USA). Emissions were recorded using a sCMOS camera (pco.edge 4.2) with 512Χ500 resolution after 4Χ4 pixel binning, and 100 ms exposure. Images were acquired using Camware software (PCO, MI, USA).

To validate the ability to resolve single cells for functional imaging under the two-photon microscope at different cortical depths, we used a Movable Objective Microscope (MOM) and galvo-resonant scanner (Sutter Instruments). Two-photon excitation was performed using a Ti:Sapphire laser (MaiTai eHP DeepSee, Spectra-Physics, CA, USA) with built-in dispersion compensation. Laser intensity into the microscope was controlled using a Pockels cell (Conoptics, CT, USA) and the laser was expanded with a 1.25Χ Galilean beam expander (A254B-100 and A254B+125, Thorlabs, NJ, USA). The laser was focused on the brain using an objective with a 1.7 mm WD and 1 NA (Plan-Apochromat 20Χ, Zeiss). Fluorescence emissions from the brain were reflected into the emissions path by a FF735Di-02 dichroic mirror (Semrock), filtered with an ET500lp long pass filter (Chroma), and then split by a T565lpxr dichroic mirror (Chroma) into two GaAsP PMTs (H10770PA-40, Hamamatsu, NJ, USA) with ET525/50m-2p (Chroma) and ET605/70m-2p (Chroma) filters for detection of green and red emissions, respectively. The two-photon microscope was controlled using ScanImage 2017 (Vidrio Technologies, VA, USA) and images were acquired at 512Χ512 resolution without bi-directional scanning.

#### Calcium imaging data analysis

Two-photon data were motion corrected for x-y displacements by rigid body registration using the moco toolbox [5] in ImageJ (NIH). Motion-corrected frames were tophat filtered across time to compensate for whole frame changes in brightness. ROIs are manually selected, and neuropil signal is removed from each ROI’s fluorescence signal as described previously [6,7]. ΔF/F was calculated for each cell using the 10^th^ percentile as the baseline. For widefield imaging data, ΔF/F for each pixel was calculated by setting the baseline to the 10^th^ percentile value for each pixel across time.

#### Statistics

Microsoft Excel 2016 and GraphPad Prism 7.01 (GraphPad Software, CA, USA) were used for data analysis and graph generation. Animal group sizes were chosen based on preliminary data that suggested a large effect size. One animal from each group was excluded from analysis after necropsy due to failed sinus injections. Final group sizes are: AAV9 n-SIM at P1 (n=6) and P4 (n=5), AAV1 n-SIM at P1 (n=4), AAV5 n-SIM at P1 (n=4), AAV-PHPeB n-SIM at P1 (n=4), temporal vein injection at P1 (n=4).

## Results and Discussion

### AAV9 n-SIM yields robust whole-brain expression as early as P5

To evaluate transduction of neurons in the mouse brain, we investigated transgene expression after n-SIM before the closure of the BBB. We examined n-SIM at P0-P1 of AAV9 expressing GCaMP6 under the control of the synapsin promoter (AAV9-syn-GCaMP6s) (Figure 1A) and observed widespread expression of GCaMP at P5 (Figure 1B,C) and persisting into adulthood (Figure 1D,E). With n-SIM, we observed labeling of 52 +/−12%, 50 +/−6%, and 51 +/−14% of cortical neurons at P6, P9, P14 respectively (Figure 1H, P6, n = 5; P9, n = 2; P14, n=5), and 66 +/−16%, 49 +/−5%, 54 +/−4% of thalamic neurons (Figure 1H, P6=2, P9, n = 2; P14, n =4).

**Figure 1.**
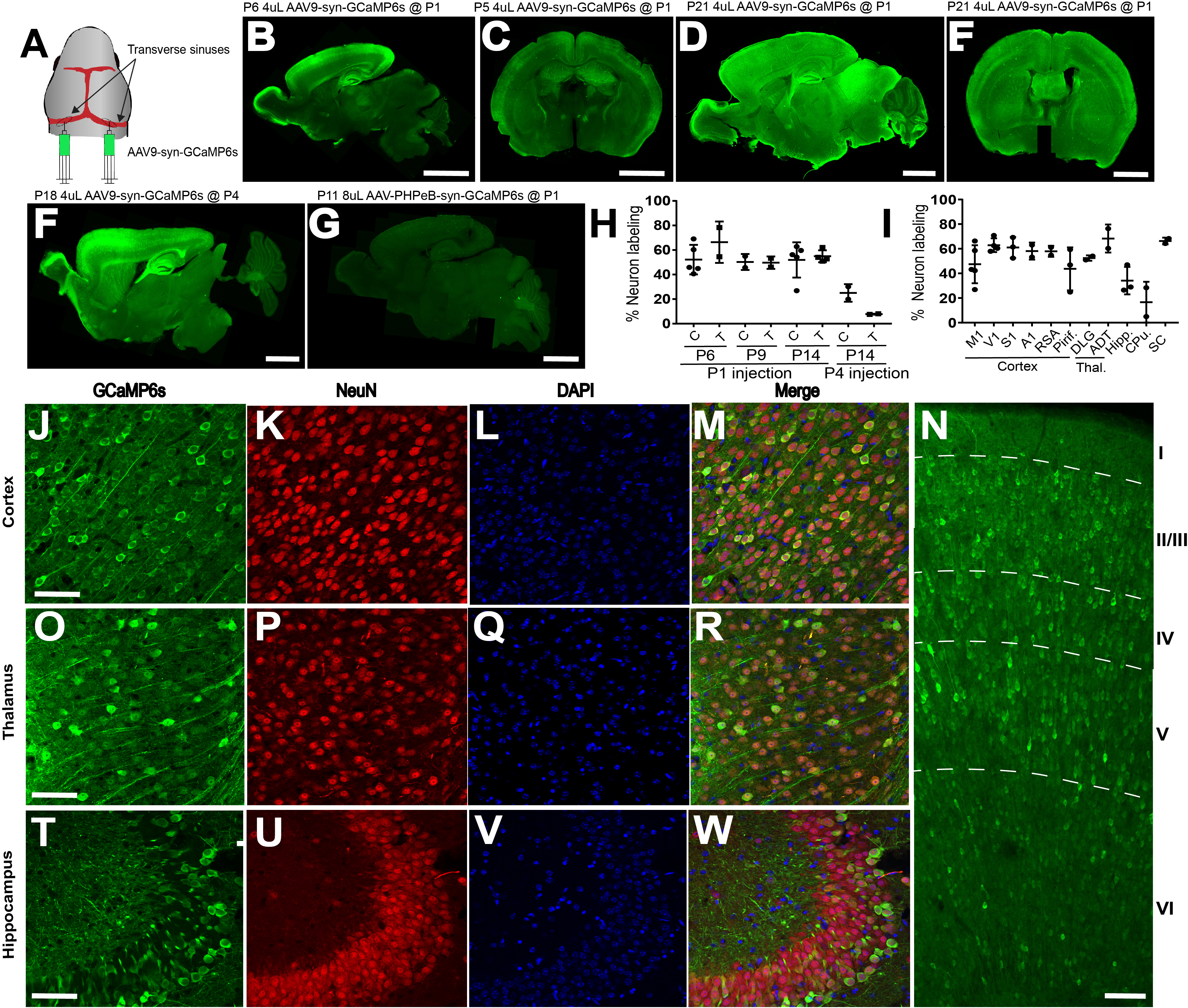
AAV9 n-SIM at P0-P1, but not at P4, leads to widespread neuronal transduction in the neonatal mouse brain. **A**: Schematic showing sites of viral injection at P1. **B,C**: Example sagittal and coronal sections of a P6 and P5 mouse brains, respectively, showing widespread and even expression of GCaMP6s across the cortex and several other brain regions including hippocampus, midbrain, and thalamus. Scale bar= 2mm. Exposure time: 1000ms. Brightness levels: **B,** 73-1200; **C,** 0-4095. **D-E**: Example sagittal and coronal sections at P21 showing that expression of GCaMP6s at P21 becomes significantly brighter relative to earlier ages. Exposure time: 1000ms. Brightness levels=0-4095. **F:** Sagittal section of P21 brain injected at P4 with 4μL of AAV9 (1×10 ^13^ vg/mL. Exposure time: 1000ms. Brightness levels=73-2000. **G:** Sagittal section of P11 brain injected with 8μL of AAV-PHPeB-syn-GCaMP6s (1×10^13^ vg/mL). Exposure time: 1000ms. Brightness levels=73-500ms. **H.** Quantification of cortical and thalamic neuron labeling at P6, P9, and P14 after P1 (P6, n=5; P9, n=2; P14, n=5) or P4 (P14, n=2) sinus injections, each data point on the plot represents an individual brain. Horizontal lines represent the mean, and vertical lines represent the standard deviation. C=cortex, T=thalamus. **I**. Quantification of neuron labeling at P14 in different cortical and thalamic regions, in addition to hippocampus and striatum. M1= motor cortex (n=5), V1= visual cortex (n=4), S1= somatosensory cortex (n=3), A1= auditory cortex (n=2), RSA= retrosplenial cortex (n=2), Piri= piriform cortex (n=3), DLG= dorsolateral geniculate nucleus (n=2), ADT= anterodorsal thalamic nucleus (n=1), Hipp= hippocampus (n=3), CPu= caudate and putamen (n=2), SC= superior colliculus (n=2). Confocal images showing GCaMP6s expression in mouse cortex and thalamus (**J-W**) at P14 after transverse sinus injection of 4μL of 1×10^13^ vg/mL AAV9-syn-GCaMP6s at P1. Panels **J**, **O**, **T** show abundant GCaMP6s expression in cortex, thalamus, and hippocampus, and show localization with both NeuN and DAPI (**M,R,W**). Scale bar= 20μm. **N**. Confocal image showing dense and widespread expression across all cortical layers at P14. Scale bar is 40μm.

Next, we investigated GCaMP expression in different cortical subregions. We found that GCaMP expression was robust in all cortical regions examined (Figure 1I; J-M; P14: M1: 47 +/−15%, n=5; V1: 62 +/−5%, n=4; S1: 61 +/−8%, n=3; A1: 58 +/−6%, n=2; Retrosplenial: 57 +/−4, n=2; Piriform: 43 +/−17%, n=3), and across all cortical layers (Figure 1R, P14, layer 2/3: 52 +/−17%; layer 4: 63 +/−13%, layer 5: 61 +/−12%, layer 6: 53 +/−17%; n=5).

We also observed robust expression in different thalamic subregions (Figure 1I; O-R; P14: DLG: 52 +/−2%; n=2, Anterodorsal thalamic nucleus: 60%, n=1), hippocampus (Figure 1I; T-W; P14: 43 +/−28%, n=3), striatum (Figure 1I; 16 +/−16%, n=2), and superior colliculus (Figure 1I; 66 +/−2%, n=2).

Overall, our results compare favorably to the sparse transfection levels reported previously using temporal vein injections of AAV9 at P1. For example, Foust et al. found 15% and 18% transfection rate in cortex at P11 and P20, respectively, after temporal vein injection at P1 (Foust et al., 2009). However, the difference in our observations could be due to the AAV9 vector that expresses GFP under the control of chicken B-actin hybrid (CB) promoter used by Foust et al. Therefore, for a more direct comparison, we injected an AAV9 vector that expresses GCaMP6s under the synapsin promoter in the temporal vein at P1 and observed expression at P14 and P21. We observed drastically weaker, slower, and non-uniform GCaMP signal relative to AAV9 delivery through the transverse sinuses. In our hands, temporal vein injections do not yield robust and uniform GCaMP expression until P21 (Supplementary Figure 2A,B,F).

To investigate whether sinus injections of AAV9 later than P1 yields similar levels of expression to injections at P1/P0, we performed injections at P4 and observed significantly lower expression levels across the brain at P14 (Figure 1F,H; P18, cortex: 25 +/−7%; thalamus: 8 +/−0%, n=2), likely due to the rapid maturation of the BBB.

To compare the efficacy of n-SIM using AAV9 relative to other serotypes, we tested n-SIM using AAV-PHPeB, AAV5, and AAV1. All three serotypes yielded relatively low expression levels even when up to double the volume (8μL) of virus was injected (Figure 1G, Supplementary Figure 2C-F). Therefore, to our knowledge, AAV9 n-SIM at P0-P1 is the only method to yield robust and widespread transgene expression in the neonatal brain without relying on transgenic driver lines.

Finally, the minimally invasive injection procedure resulted in overall health of sinus-injected animals that was comparable to non-injected controls as measured by weight-gain of the animals (P7-P9 sinus injected at P1: 5.1 +/−0.84g, n=8; P7-P9 non-injected: 5 +/−1g, n=5). Sinus injected animals survived for as long as we have observed (P63, n=3), with no evidence of infection or rejection by the dam.

### AAV9 n-SIM at P1 enables efficient labeling of different cell populations in the same brain during the first postnatal week

With the unique ability to efficiently label and manipulate brain cells very early in postnatal development, our method brings us closer to understanding how different subtypes of neurons (e.g. interneurons) are integrated into cortical circuits. Little is known about how different subtypes of interneurons are functionally integrated into circuits in the developing cortex. Moreover, sparse interneuron populations (e.g. VIP interneurons, which account for less than 2% of cortical neurons) are not easily accessible with electrophysiology and are difficult to label and manipulate during the first postnatal week [9]. Furthermore, interneuron activity is critical for the proper development of cortical circuitry [10], but little is known about their relationship to overall cortical dynamics, or how their early disruption may affect cortical function. In summary, to study interneuron activity in relation to surrounding pyramidal neurons and to be able to conditionally manipulate them as early as P5, is difficult to accomplish using present labeling methods (e.g. cortical injection) as interneurons do not finish migrating into the cortical plate until P7 [10]. Our method circumvents the methodological challenges of labeling sparse neuronal populations, while preserving the possibility of combining functional imaging with genetic labeling. Labeling using our method results in sufficiently bright GCaMP expression to perform two-photon calcium imaging of layer 2/3 excitatory and inhibitory (Nkx2.1, somatostatin (SOM), VIP) neurons simultaneously as early as P4 (Figure 2A-I) and persisting into adulthood [11]. In addition to imaging layer 2/3, labeling through n-SIM also enabled us to image deep cortical layers as early as P9 (Figure 2H,I). Time-series traces (Figure 2C,E,G,I) show that interneuron subtypes in V1 displayed spontaneous activity as shown by their calcium transients. Therefore, our method is viable to use for labeling and manipulating excitatory and inhibitory neurons across the whole brain during the first postnatal week.

**Figure 2.**
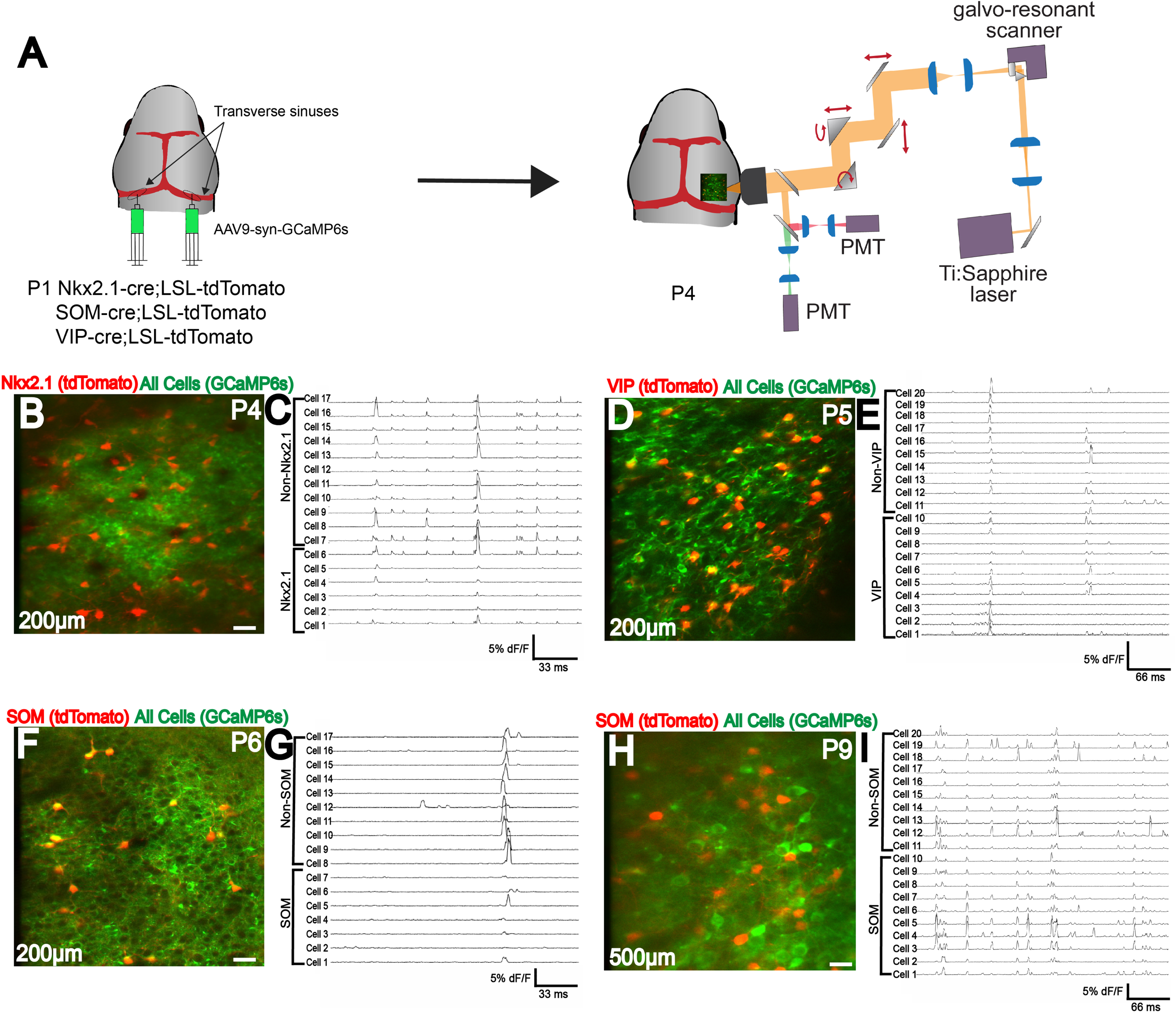
Efficient labeling of multiple cell types in neonatal cortex using n-SIM. **A**. Schematic showing viral injection (AAV9-syn-GCaMP6s) into Nkx2.1;LSL-tdTomato, or SOM-cre;LSL-tdTomato, or VIP-cre;LSL-tdTomato to label excitatory and inhibitory neuron populations. 2-photon imaging setup is shown. **B-I**. Simultaneous two-photon imaging of Nkx2.1, SOM, and VIP interneurons in V1 layers 2/3 along with surrounding pyramidal neurons as early as P4, showing efficient labeling of all interneuron subtypes. Traces adjacent to images show that all interneuron subtypes display calcium transients. **H,I**. Deep-layer two-photon imaging of SOM interneurons and surrounding pyramidal neurons at P9, showing efficient labeling of deep cortical layers early in development. Note that for illustration purposes, only a subset of neurons have traces displayed in this figure. Scale bar is 20μm.

### Simultaneous whole-brain expression of two constructs

In addition to the expression of single transgenes, n-SIM provides the ability to achieve whole-brain expression of multiple constructs simultaneously. For instance, we expressed both GCaMP6s and jRCaMP1b across the brain in specific cell types using Cre recombinase in VIP interneurons. To do this, we co-injected a total volume of 4μL of AAV9-CAG-flex-GCaMP6s and AAV9-syn-jRCaMP1b (1:1 ratio) at P1 (Figure 3A). With this preparation, we observed wide spread expression across the cortical mantle, suitable for macroscopic calcium imaging of VIP interneurons (with GCaMP6) and all neurons (with RCaMP) at P10 (Figure 3B-E). This allows us to visualize and distinguish calcium dynamics from two neuronal populations simultaneously and separately under the single-photon mesoscope. Using this approach, we are currently performing simultaneous imaging of neural populations across the whole brain by exciting GCaMP6s at 465nm and jRCaMP1b at 565nm. This allows us to measure and compare the spatiotemporal dynamics of excitatory and inhibitory neurons cortex-wide throughout development, which is currently inaccessible by any other methodology to the best of our knowledge. Therefore, this novel method will enable us to study how interneurons are integrated into functional circuits in the cortex, or how their early disruption may affect cortical function.

**Figure 3.**
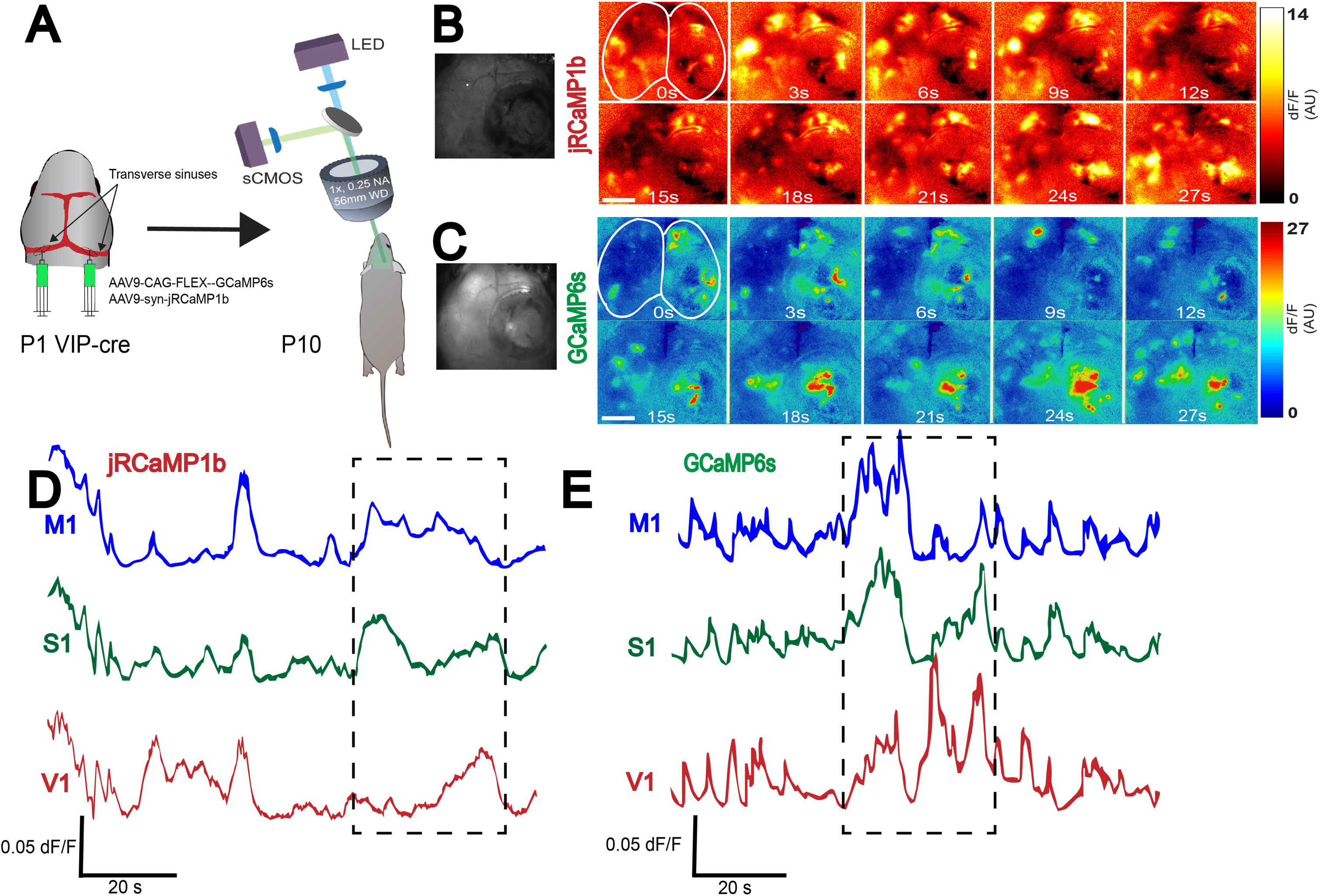
Whole-brain expression of multiple viral constructs in neonates. **A.** Schematic showing co-injection of two viruses (AAV9-syn-jRCaMP1b and AAV9-CAG-flex-GCaMP6s) into the transverse sinuses of VIP-cre mice at P0-P1, and widefield imaging of two neural populations in the same brain at P10. Widefield imaging setup is shown in simplified schematic (see Methods section for details). **B,C.** Montage of neural activity across cortex imaged using the widefield mesoscope. All neurons are labeled using jRCaMP1b, and only VIP interneurons are labeled using GCaMP6s. **D,E**. Traces represent time-series of spontaneous activity measured by calcium transients from motor cortex (M1), somatosensory cortex (S1), and visual cortex (V1). Boxed area of traces in D and E are shown as a montage in **B** and **C**, respectively. Scale bar is 2μm.

### Multi-species compatibility of n-SIM

Another advantage of the n-SIM is its compatibility to work in species other than mice that lack the wide array of genetic tools, such as rats. We test n-SIM at P1 in Long Evans rats, and we are able to achieve robust whole-brain expression as early as P6 using as little as 4μL of virus (Figure 4). The GCaMP signal had comparable brightness and activity-dependent change in fluorescence (∆F/F) to mice at a similar age under the same injection conditions (Figure 4A-C). Using our method, we are able to detect and quantify spontaneous neural activity during the first week of postnatal development using both widefield (Figure 4B,C) and 2-photon calcium imaging (Figure 4D) from across cortex. Immunostaining of sinus injected rats shows robust expression of GCaMP6 across the whole brain as early as P6 as shown by confocal (Figure 4E) and epiflourescence images (Figure 4F).

**Figure 4.**
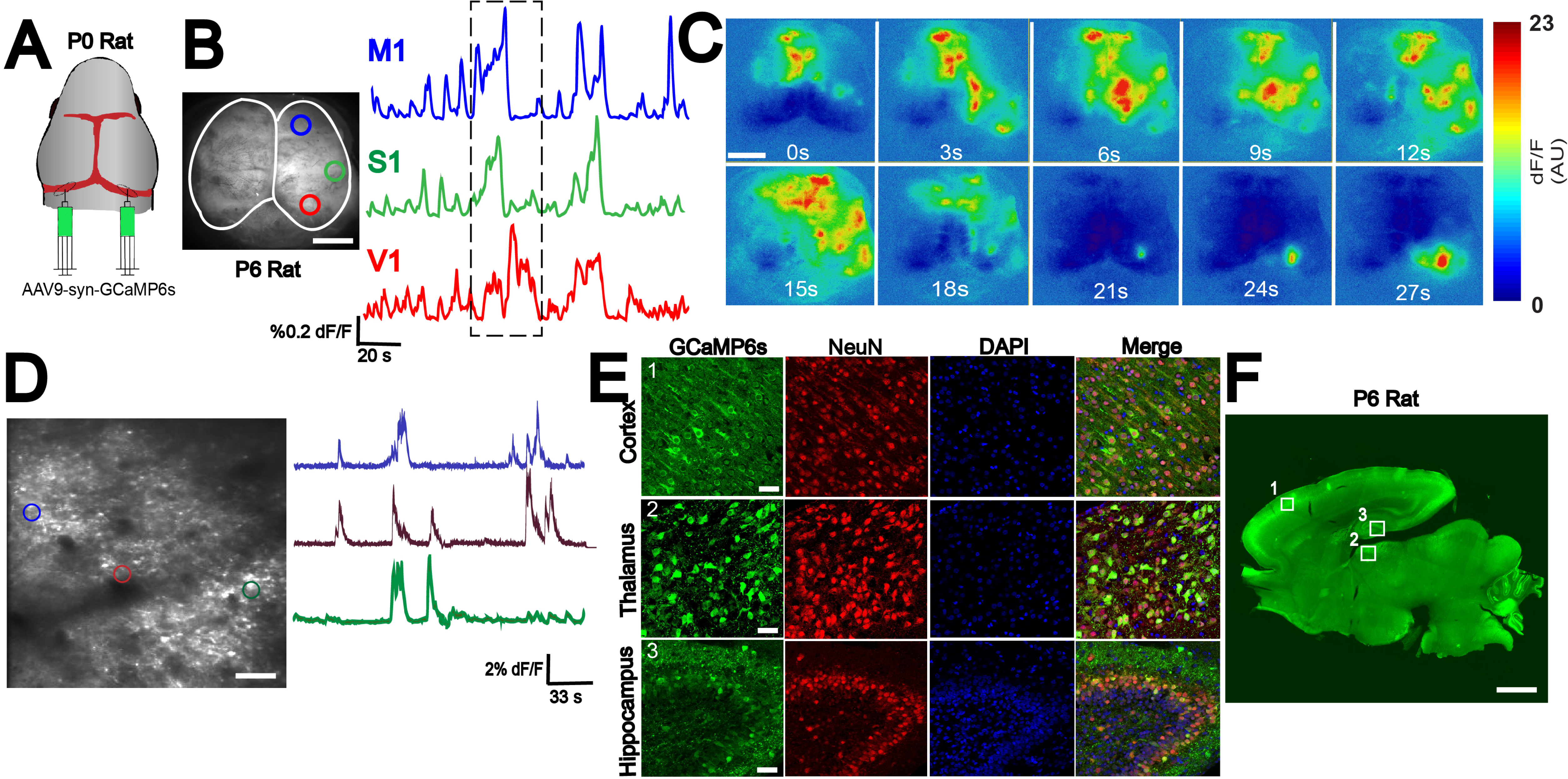
n-SIM compatibility in multiple species. **A**. n-SIM using 4μL of AAV9-μL of AAV9-L of AAV9-syn-GCaMP6s (1×10^13^ vg/mL) at P0-P1 in rats yields whole-brain expression of GCaMP at P6 with strong functional signal across cross cortex as shown by widefield imaging (**B**). Traces represent time-series of spontaneous activity measured by calcium transients from motor cortex (M1), somatosensory cortex (S1), and visual cortex (V1). **C**. Boxed area of traces in (A) shown in a montage. Scale bar is 4μL of AAV9-μL of AAV9-m. **D**. Two-photon imaging of spontaneous cortical activity in V1 from a P6 rat with sample traces from individual neurons marked with color-matched circles. Scale bar is 50μL of AAV9-m. **E.** Confocal images from boxed areas in F showing dense labeling of neurons in cortex, thalamus, and hippocampus at P6. Scale bar is 50μL of AAV9-m. **F**. Sagittal section of a P6 rat brain showing widespread rostral-caudal expression of GCaMP6 across the whole brain. Scale bar is 2mm. Exposure time: 6000ms. Brightness levels=73-2500.

Labeling with our method provides the opportunity to answer a wide variety of fundamental questions in neuroscience that were previously impossible to approach, such as inter-species comparisons of cortical development. Furthermore, the rules that govern cortical development prior to onset of sensation in different species are not clear. However, it is evident that spontaneous neuronal activity prior to eye opening instructs cortical development [12]. Using our method, we have already begun examining the patterns of spontaneous cortical activity in developing rats prior to eye opening (~P12) and making direct comparisons to mice.

In summary, our results demonstrate a novel method to achieve robust whole-brain expression of transgenes in neonates and enables the labeling and imaging two neuronal populations simultaneously across development. Our method is also applicable for use in different mammalian species and will help us unravel the principles underlying the development of functional organization of cortex. Our study provides a proof of concept of a powerful tool that may also be of clinical relevance.

## Acknowledgments

This research was supported by NIH F32 EY028869-01A1, R01 EY015788, R01 EY023105, U01 NS094358, P30 EY026878, R01 MH111424. We thank Yueyi Zhang for help with animal breeding and maintenance, Xinxin Ge for help with data analysis, R. Todd Constable, Michael Higley, Daniel Barson, and Jess Cardin for help with building the microscope. We also thank Drs. Evelyn Lake and Alicja Puscian for helpful comments on the manuscript. The authors declare no competing financial interests.

---

**Supplementary Figure 1.**
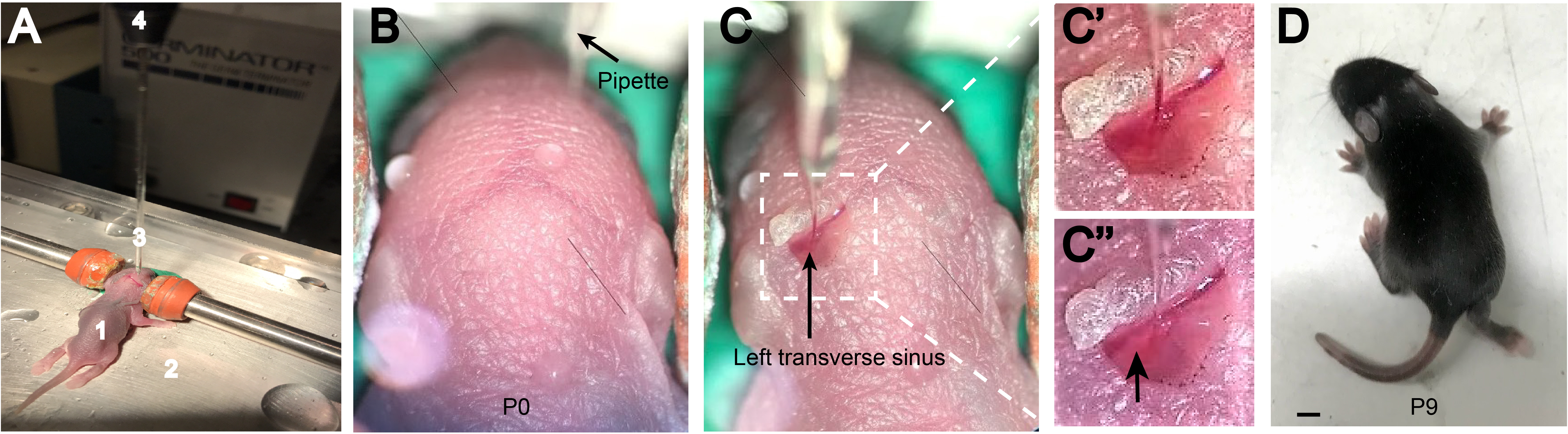
Neonatal transverse sinus injection method. **A.** Injection set-up showing P0 pup (1) on ice-cold plate (2) with glass pipette (3) and nanoject (4) positioned above site of injection. **B.** View through the dissection microscope showing dorsal side of the head before any incisions are made. **C.** After incision is made to expose the sinuses, the glass pipette is lowered through the skull such that the tip of pipette is within the lumen of the sinus. Once we begin the viral injection (**C**’), the correct depth of injection is verified by visualizing the viral solution plume within the blood stream of the sinus (Arrowhead in **C**”). **D.** P9 sinus-injected mouse showing good would healing around the site of injection. Scale bar= 2mm.

**Supplementary Figure 2.**
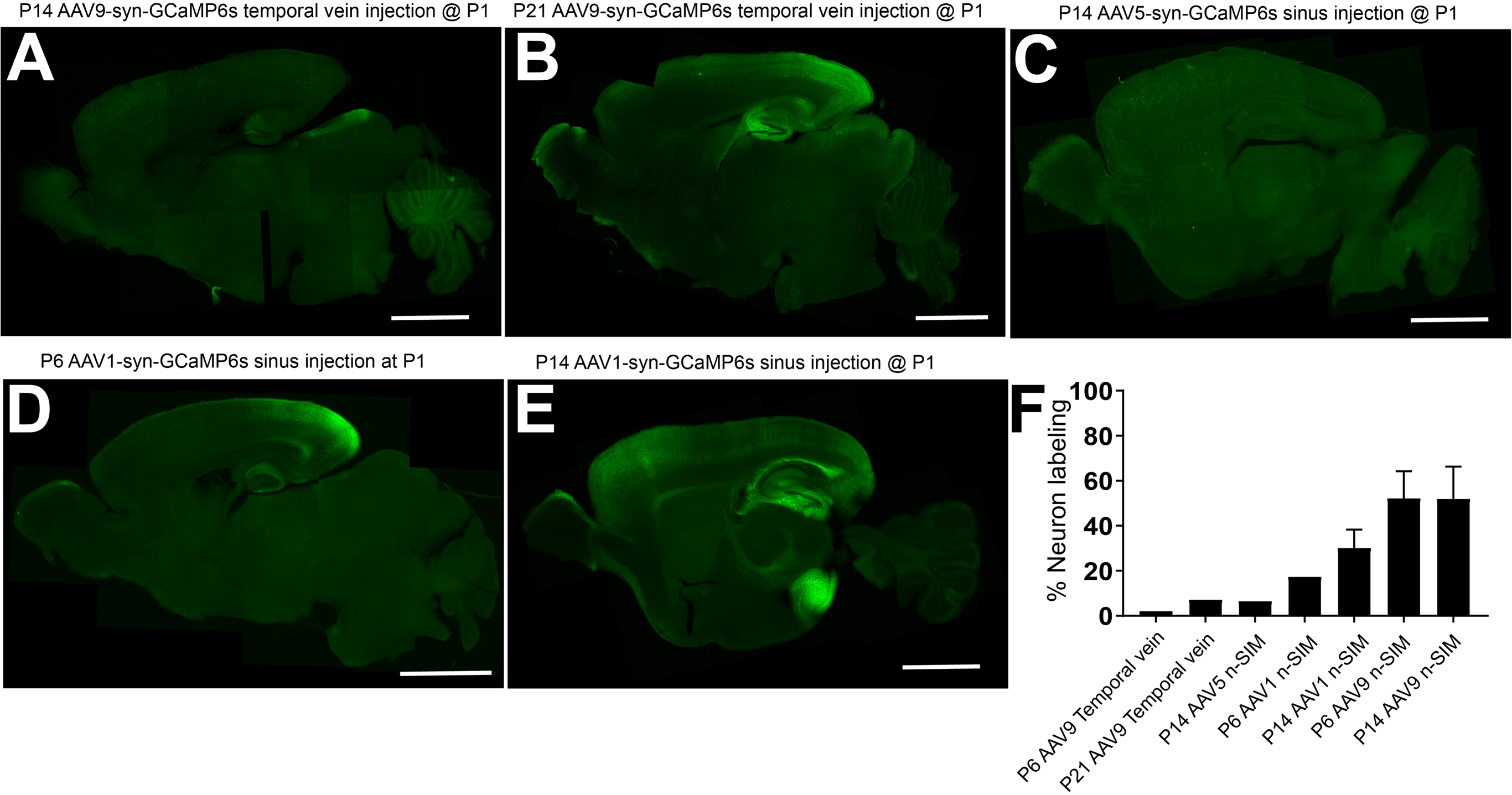
Comparison of AAV9 n-SIM to other methods. **A,B**. Sagittal sections of P14 and P21 mouse brains with bilateral temporal vein injections of AAV9-syn-GCaMP6s at P1. **A**: Exposure time= 2000ms. Brightness levels=73-2500. **B**: Exposure time= 1000ms. Brightness levels=73-2500. **C.** Sagittal sections of P14 mouse brains with AAV5 n-SIM at P1. Exposure time= 1000ms. Brightness levels=73-2500. **D,E.** Sagittal sections of P6 and P14 mouse brains labeled with GCaMP6s using AAV1 n-SIM at P1. Exposure time= 2000ms. Brightness levels=73-2500. **F**: Quantification of neuron labeling at P6, P14, and P21 using AAV9 temporal vein injections at P1, AAV5 n-SIM at P1, and AAV1 n-SIM at P1. Note comparison to AAV9 n-SIM at P1 (refer to Figure 1 H). Scale bars are 2mm.

